# Germline-soma Supply Mitochondria for mtDNA Inheritance in Mouse Oogenesis

**DOI:** 10.1101/2022.01.11.475948

**Authors:** Hongying Sha, Zhao Ye, Zhen Ye, Sanbao Shi, Jianxin Pan, Xi Dong, Yao Zhao

**Affiliations:** Reproductive Medicine Center, Zhongshan Hospital, Shanghai Medical College, Fudan University; Shanghai 200040, China; Hypothalamus-pituitary Research Center, Department of Neurosurgery, Huashan Hospital, Shanghai Medical College, Fudan University; Shanghai 200040, China; State Key Laboratory of Medical Neurobiology, Institutes of Brain Science, Shanghai Medical College, Fudan University; Shanghai, 200040, China

## Abstract

Maternal transmission paradigm of mtDNA remains controversial in mammalian oogenesis. Germline-soma-to-oocyte communication by numerous transzonal nanotubes (TZTs) reminds whether intercellular mitochondrial transfer is associated with maternal inheritance. Here, we found that mouse oocytes egocentrically receive mitochondria via TZTs, which projected from germline-soma, to achieve 10^5^ copies, instead of *de novo* synthesis of mtDNA subpopulation in growing oocytes. *De novo* assembled TZTs amongst germline-soma and oocytes accumulated mtDNA amounts of the oocytes *in vitro*. However, mitochondrial supplement from germline-soma gradually diminished along with oocyte growth and was terminated by meiosis resumption, in line with a decrease in the proportion of germline-soma with thriving mtDNA replication and FSH capture capability. Thus, germline-soma-to-oocyte mitochondrial transfer is responsible for mammalian mtDNA inheritance as well as oogenesis and aging.

**One-sentence summaries:** The maternal mtDNA transmission accompanied by oogenesis is inseparable from germline-soma mitochondrial transport.

## Main Text

Maternal transmission paradigm of mtDNA remains controversial in mammalian oogenesis and has raised a number of important considerations *(1, 2, 3)*. Mathematically, 10-100 copies of mtDNA in a primordial germ cell require 9-14 times of replication for each genetic molecule to achieve 10^5^ copies during mammalian oogenesis *(4, 5)*. MtDNA is particularly vulnerable *in vivo* due to the lack of protective histone proteins and accurate repair mechanisms, which compromises their function and leads to mutation. While germline selection would purify severe mtDNA mutant *(6)*, the paucity of mutation in maternal pedigrees suggested an unexpected mechanism for mitochondrial biogenesis, which provided stable inheritance of maternal mitochondria during oogenesis. The intercellular transfer of mitochondria, with the discovery of tunneling nanotubes, reveals a novel biological principle in the development and differentiation of multicellular organisms *(7-13)*. Mouse oocytes grow in a microenvironment, in which germline-soma, especially cumulus cells, elaborate numerous transzonal nanotubes (TZTs) to reach the surface of oocytes for mutual contact *(14,15)*. Here, we demonstrated whether and how mitochondria transport between cumulus cells and oocytes is coupled with mtDNA inheritance in mammalian oogenesis.

We found that almost all mitochondria are short and round in the growing mouse oocytes (Fig. S1). Co-labeling with EdU and anti-DNA antibody showed the absence of EdU signals in oocyte cytoplasm, while intensive EdU signals, indicating mtDNA replication, occurred in cumulus cells (Fig. 1A, Fig. S2). Twinkle, known as the helicase of mtDNA replication in mammals, was absent from the oocyte cytoplasm (Fig. 1B). MFF, which only governs midzone fission and leads to mitochondrial proliferation *(16)*, appeared deficient as well (Fig. 1C). However, both twinkle-positive and MFF-positive cumulus cells located in COCs (Fig. 1B, C). Contrarily, FIS, which regulates peripheral fission *(16)*, present in whole COCs (Fig. S3). Besides, we found that cumulus cells directly contacted the membranes of oocytes by stretching TZTs (Fig. 1D, E), some of which contained mitochondria (Fig. 1F). Notably, *in vitro* maturation (IVM) failed to accumulate mtDNA copies in oocytes (Fig. 1G, H, Tables. S2), as its insufficiency in producing oocytes with quality comparable to the oocytes matured *in vivo (17,18)*. Together, these data indicated that *de novo* synthesis of mtDNA is absent in mouse growing oocytes, and mtDNA copies in oocytes no longer increase when cumulus cells were disaggregated during maturation *in vitro*.

**Fig. 1.**
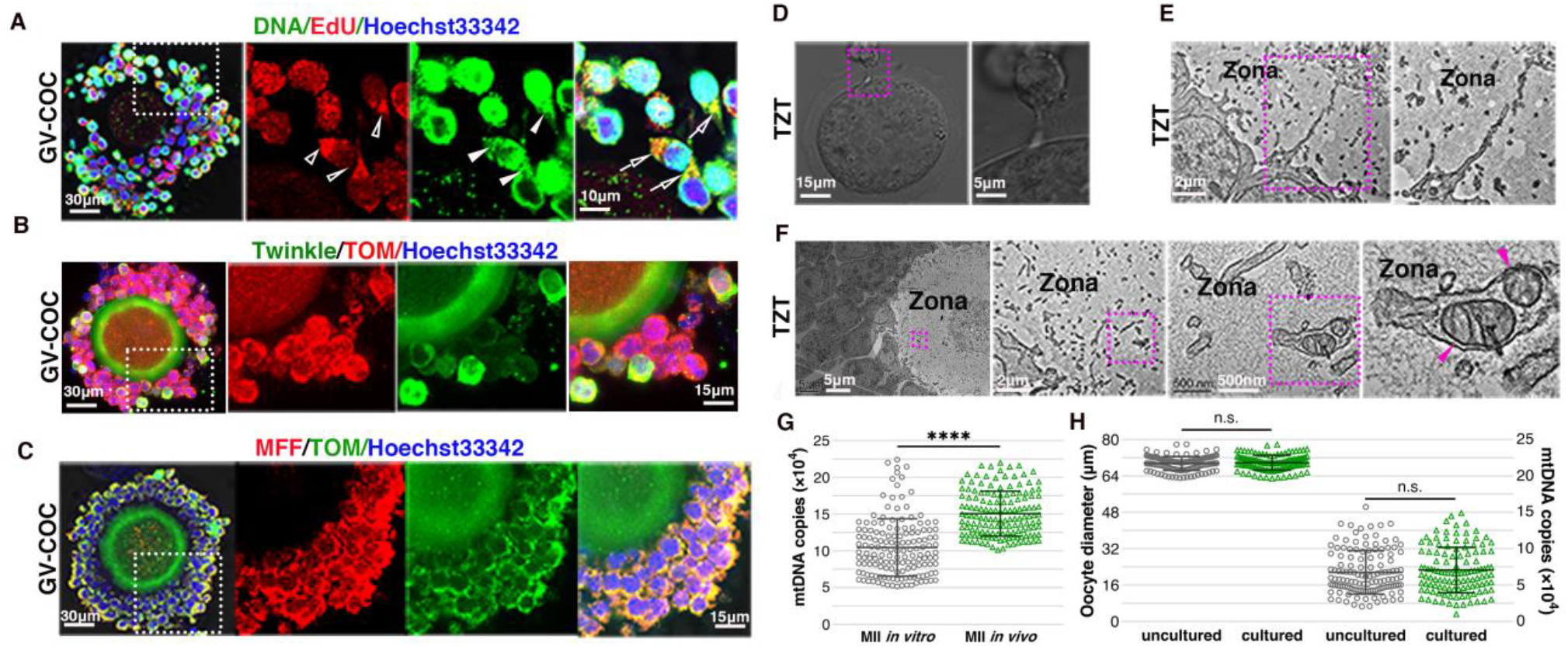
Mitochondrial biogenesis was absent in growing mouse oocytes. **(A)** Representative images of GV-COCs labeled with EdU (red) and anti-DNA antibody (green). Hollow arrowheads indicate EdU, solid arrowheads indicate mtDNA, arrows indicate co-location of EdU and mtDNA. **(B)** Representative images of GV-COCs labeled with Twinkle (green) and TOM (red). **(C)** Representative images of GV-COCs labeled with TOM (green) and MFF (red). **(D)** Architecture of a TZT between cumulus and oocyte. **(E)** Intact ultrastructure of a TZT between cumulus and oocytes detected by TEM of 70 nm section. **(F)** A discontinuous TZT, which is disrupted by TEM analysis, contained two mitochondria. For boxed areas, higher magnification images were shown. Pink arrowheads indicate mitochondria in TZT. **(G)** MtDNA copies of MII oocytes from *in vivo* and *in vitro*, respectively. **(H)** MtDNA copies of Oocytes cultured 48 hours *in vitro* and uncultured oocytes, respectively. *In vitro* culture of oocytes was performed with contactless co-culture with cumulus cells. Each data point represents an individual oocyte mtDNA copies. Data are shown as mean ± SD. n.s., not significant, ****P < 0.0001.

Thus, we sought to explore whether cumulus cells transfer mitochondria into oocytes for mtDNA accumulation in mouse oogenesis. We first stained COCs at germinal vesicle (GV-COCs) and at the first meiosis (MI-COCs) with Nonyl Acridine Orange (NAO) using a spatiotemporal-staining method, by which NAO only stain cumulus cells while the oocytes remained uncolored in COCs through controlling staining time (Fig. S4A), and subsequently cultured for 1 and 5 h in a defined pre-IVM medium with meiotic inhibitor (10μM cilostamide). We found that prolonged COCs culture drastically enhanced NAO intensity in the GV oocytes (Fig. 2A, B, C, Fig. S5∼7, Table. S3, P<0.0001). But the NAO intensity in the oocytes from 5 h group gradually declined as the diameter increased, while 1 h group maintained a constant low level (Fig. 2D). ΔNAO intensity between the two groups appears to be negatively correlated with oocyte diameter (Fig. 2E).

**Fig. 2.**
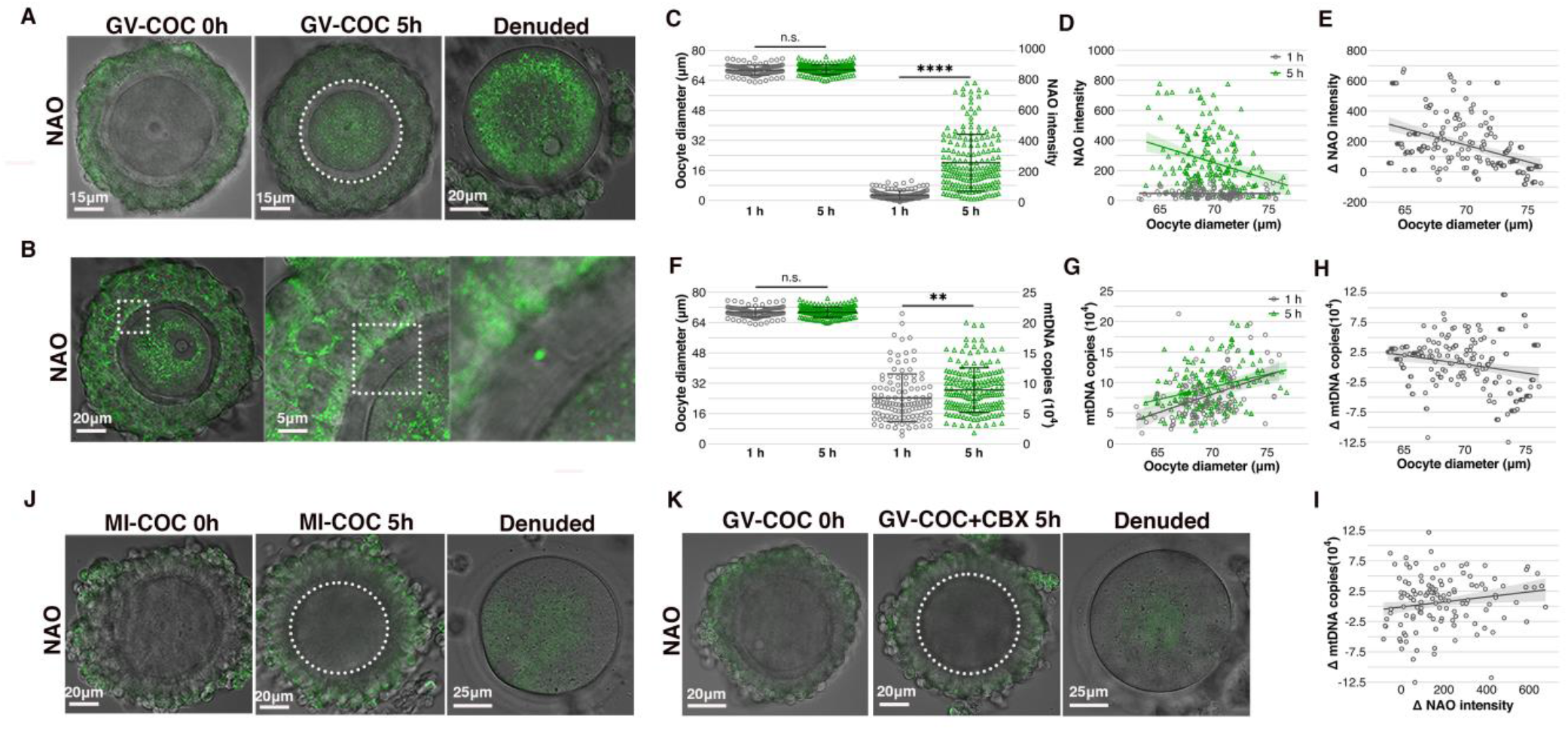
Mitochondria enriched in mouse oocytes during the COCs maturation *in vitro*. **(A)** Representative images of GV-COCs labeled with NAO. Left, pre-cultured; middle, cultured 5h; right, oocyte from circled areas. **(B)** Representative confocal image showed the localization of NAO signal on *zona pellucida* after GV-COCs cultured 5h *in vitro*. For boxed areas, higher magnification images are shown. **(C)** Quantification of NAO intensity in the GV oocytes after COCs cultured 1h and 5h, respectively. Each data point represents an individual oocyte NAO intensity. **(D)** The trend of NAO intensity with the increase of oocyte diameter from 1h and 5h groups. Each data point represents an individual oocyte NAO intensity. (t-test, P_slope_ = 0.0004). **(E)** The trend of ΔNAO intensity with the increase of oocytes diameter (Pearson correlation, r = - 0.4415, P < 0.0001). ΔNAO intensity denotes NAO value in the GV oocytes from COCs 5h minus that from COC 1h. Each data point represents increased NAO amount of a simulated individual cell. **(F)** Quantification of mtDNA copies in the oocytes from 1h and 5h *groups*, respectively. Each data point represents an individual oocyte mtDNA copies. **(G)** The trend of mtDNA copies with the diameter increase of oocytes from 1h and 5h group. Each data point represents an individual oocyte mtDNA copies. (t-test, P_slope_=0.2023, P_intercept_=0.0017). **(H)** The trend of ΔmtDNA copies with the diameter increase of oocytes (Pearson correlation, r = -0.2368, P = 0.0012). ΔmtDNA copies denotes mtDNA copies in the GV oocytes from COCs 5h minus that from COC 1h. Each data point represents increased mtDNA copies of a simulated individual oocyte. (t-test, P_slope_ = 0.0004) **(I)** The correlation between ΔNAO intensity and ΔmtDNA copies (Pearson correlation r = 0.1671, P = 0.0223). **(J)** Representative images of MI-COCs labeled with NAO. Left, pre-cultured; middle, cultured 5h; right, oocyte from circled areas. **(K)** Representative images of GVCOCs labeled with NAO followed CBX treatment. Left, pre-cultured; middle, cultured 5h; right, oocyte from circled areas. Background fluorescent noise in A, B, J and K were not normalized using an Offset parameter at -25.0. The lines and shaded region showed linear correlation with 95% confidence intervals. Data are shown as mean ± SD. n.s., not significant, **P < 0.01. ****P < 0.0001.

We next observed that prolonged COCs culture also increased the mtDNA copies in the GV oocytes (Fig. 2F, Table. S4, P<0.01). Different from the NAO value, the mtDNA copies in the two groups showed an upward trend with the increase of oocyte diameter, but the regression curve of 5h group always located above 1h group (Fig. 2G). ΔmtDNA copies between the two groups is also negatively correlated with oocyte diameter (Fig. 2H). However, ΔNAO intensity is positively correlative to ΔmtDNA copies in the oocyte (Fig. 2I, P<0.05). Significantly, we found meiosis resumption or gap junction blocker terminated the NAO movements, as no NAO signal present in MI-group and the blocker treated GV-COCs group (Fig. 2J, K, Fig. S4B, C, Table. S5), correspondent with the fact that both luteinizing hormone and gap junction blocker disrupt TZT junction between cumulus cells and oocytes *(19)*. Therefore, the above results suggested that the fluorescent signal occurred in oocytes may be NAO probe-bound mitochondria (mtDNA) moved from cumulus cells via TZTs.

Considering the fluorescence signal occurred in the GV oocytes may be the leakage of NAO dye during COCs staining, TZTs were formed *de novo* by mixing the NAO-stained cumulus cells and unstained oocytes (GV and MI) together to generate reconstructed COCs (re-COCs) in a pre-IVM medium containing oligo hyaluronic acid (2000-8000Da) *(20)* and meiotic inhibitor (Fig. 3A, B, Fig. S8A). Cumulus cell-free GV oocytes were co-cultured as a control (Fig. S8B). After 28∼30 h incubation, positive NAO signals occurred in the GV oocytes from re-COCs group, in which NAO signal is halfway through a *de novo* nanotube (Fig. 3B, C). Phalloidin staining further showed that numerous *de novo* TZTs connect the cumulus cells with oocytes, whereas TZT absent in those *zona pellucida* where no cumulus cells adhere (Fig. 3D).

**Fig. 3.**
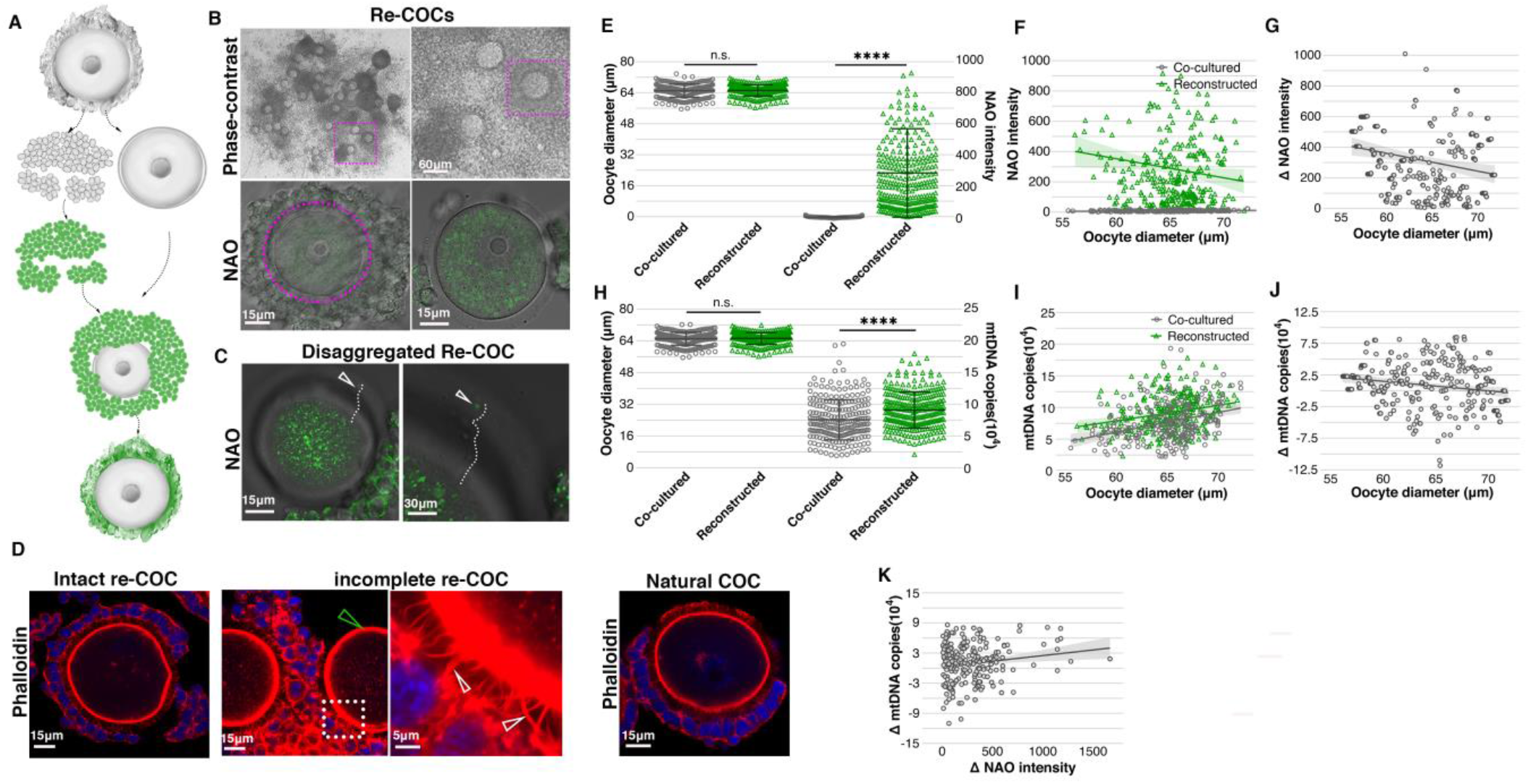
Cumulus cells assembled *de novo* TZTs to transport mitochondria into GV oocytes. **(A)** Schematic strategy of elaborating TZTs *de novo* to generate reconstructed COCs. **(B)** Representative images of re-COCs under the phase-contrast microscope (upper) and confocal microscope (lower). For boxed areas, higher magnification images are shown. The oocyte in lower right was from circled areas. **(C)** Representative confocal image showed a nanotube containing NAO signal. Dotted line simulates the direction of the nanotube. Hollow arrowheads indicated that NAO is halfway through the nanotube. **(D)** Phalloidin staining showed *de novo* TZTs connecting the oocyte with cumulus cells in re-COCs. Left, intact re-COCs; middle, incomplete re-COC; right, natural COC. For boxed areas, higher magnification images were shown. White arrowheads indicate thick TZTs. Green arrowhead indicates no TZT was established on *zona pellucida* where no cumulus cells adhere. **(E)** Quantification of NAO intensity in the GV oocytes from re-COCs and co-cultured groups, respectively. Each data point represents an individual oocyte NAO intensity. **(F)** The trend of NAO intensity with increase of the GV oocyte diameter from re-COCs and co-cultured groups. Each data point represents an individual oocyte NAO intensity. (t-test, P_slope_ = 0.0218). **(G)** The trend of ΔNAO intensity with increase of the GV oocyte diameter (Pearson correlation, r = -0.2017, P = 0.0005). ΔNAO intensity denotes fluorescence value of re-COCs minus that of co-cultured groups. Each data point represents increased NAO amount of a simulated individual oocyte. **(H)** Quantification of mtDNA copies in the GV oocytes from re-COCs and co-cultured groups, respectively. Each data point represents an individual oocyte mtDNA copies. **(I)** The trend of mtDNA copies with the increase of the GV oocyte diameter from re-COCs and co-cultured groups, respectively. Each data point represents an individual oocyte mtDNA copies. (t-test, P_slope_=0.3904, P_intercept_<0.0001). **(J)** The trend of ΔmtDNA copies with the increase of the GV oocytes diameter (Pearson correlation, r = -0.1957, p = 0.0007). ΔmtDNA copies denote mtDNA copies in the GV oocytes from re-COCs minus that from co-cultured groups. Each data point represents increased mtDNA copies of a simulated individual oocyte. **(K)** The correlation between ΔNAO intensity and ΔmtDNA copies (Pearson correlation, r = 0.1636, p = 0.0049). Re-COCs, reconstructed COCs. Background fluorescent noise in B and C were not normalized using an Offset parameter at -25.0. The lines and shaded region showed linear correlation with 95% confidence intervals. Data are shown as mean ± SD. n.s. = not significant, ****P < 0.0001.

Next, we found that *de novo* TZTs augmented the average NAO intensity in the GV oocytes from the re-COCs group (Fig. 3E, Fig. S9-12, Table. S6). Nonetheless, the NAO intensity in the re-COC oocytes gradually dropped as the oocyte diameter increased, while the co-culture group remained a constant low level (Fig. 3F). ΔNAO intensity between the two groups appeared to be negatively correlated with the oocyte diameter (Fig. 3G). *De novo* TZTs also increased the mtDNA copies in the GV oocytes from the re-COCs group (Fig. 3H, Table. S7). Different from the NAO value, the mtDNA copies in two groups showed an upward trend with the increase in oocyte diameter, but the regression curve of the re-COCs group always reside upon the co-culture group (Fig. 3I). ΔmtDNA copies between the two groups, is negatively associated with the oocyte diameter (Fig. 3J). However, a significant positive correlation occurred between the ΔNAO intensity and ΔmtDNA copies (Fig. 3K). Significantly, no NAO signal present in MI oocytes, which was reaggregated for 28∼30 h with NAO-stained cumulus cells (Fig. S8C, D, Table. S8). These results suggested that the NAO intensity in GV oocytes represent the transferred mitochondria (mtDNA) from cumulus cells, and the amount of that gradually decreased as the oocytes grew. Our results provided the mechanism why in animals and humans, mitochondrial morphology in the oocyte corresponded to that in its sister cumulus cells, in which aging oocyte and its cumulus cells contain elongated mitochondria with high-density matrix particles, while young group possesses most round mitochondria with homogenous matrix density *(21-23)*.

To reveal how the mtDNA accumulation are regulated in growing oocytes, we next explored potential links between mtDNA replication, follicle-stimulating hormone receptor (FSHr) expression and cumulus cell proliferation during follicular growth, as the maximum increment of mtDNA copies occurred at the antral follicle stage *(2, 5)*, where FSHr captures FSH to promote cumulus cell proliferation and oocyte growth *(24, 25)*. EdU incorporation showed that cumulus cells continued proliferating with a stable upward trend during COCs growth and maturation (Fig. 4A, B). In contrast to this, both twinkle and FSHr demonstrated distinct kinetics, which had a sudden rise and then dropped (Fig. 4A, B). Co-labeling showed that twinkle and FSHr were co-located in the same cumulus cells (Fig. 4C). Significantly, we found twinkle-positive cumulus cells stretch thick tubule through the *zona pellucida* to connect oocyte membrane (Fig. 4D), implying twinkle-positive and FSHr-positive cumulus cells may supply mitochondria for the mtDNA accumulation in growing oocytes. Further, GV oocytes tended to recruit twinkle/FSHr-positive cumulus cells to assemble TZTs *de novo* and generate re-COCs *in vitro* (Fig. 4E, F). Consequently, we speculate that gonadotropin hormone, such as FSH, released from hypothalamic pituitary, may regulate mtDNA replication and mitochondrial biogenesis in cumulus cells, and subsequently transport them into oocytes by the variation of FSHr level (Fig. 4G). Our data explains why a reduction in the number of TZTs associated with aging is responsible for age-related mitochondrial dysfunction and meiotic defects *(26)*.

**Fig. 4.**
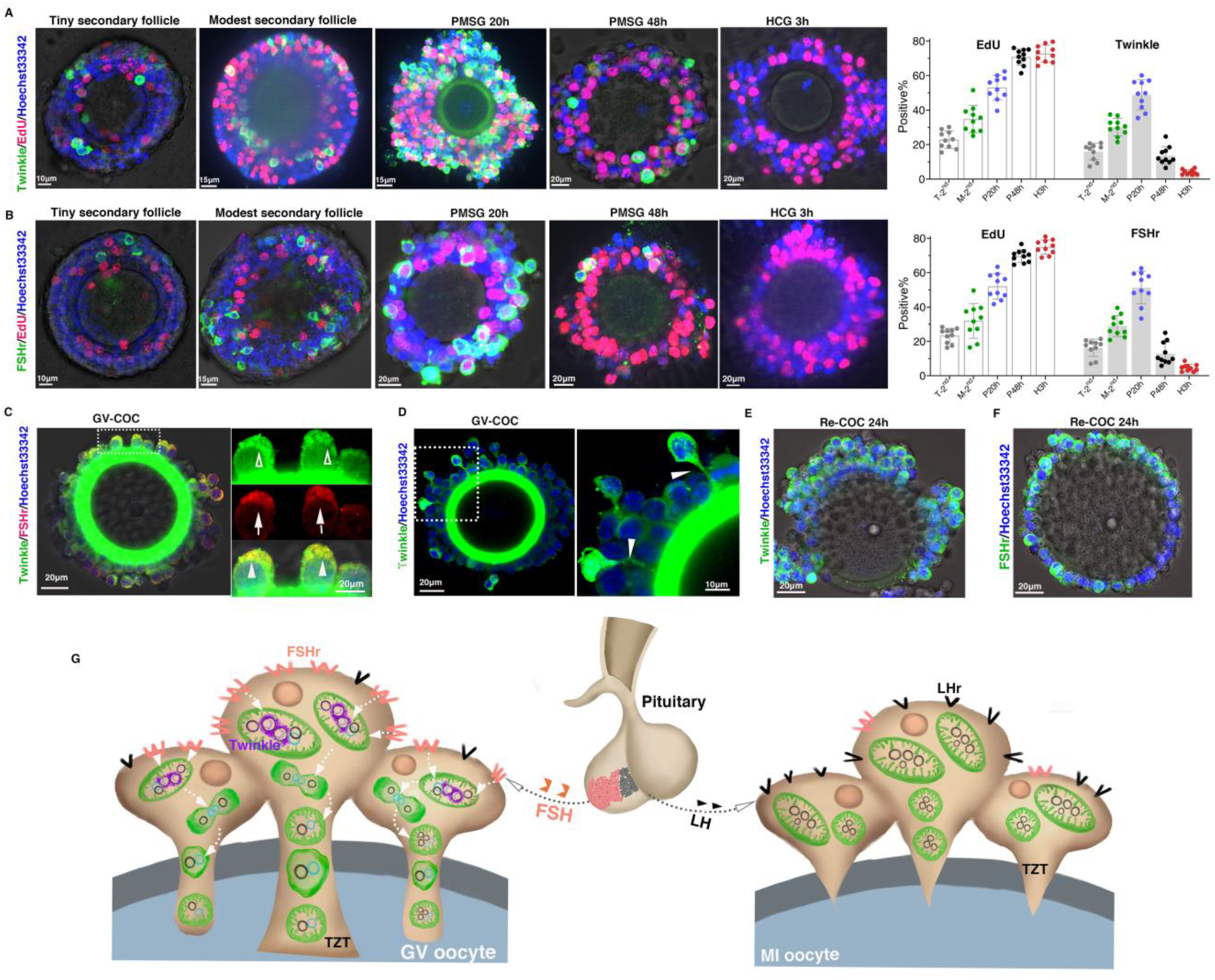
Mechanism underly when and how germline-soma transfer mitochondria into mouse oocytes. **(A)** Left, representative confocal image of the localization of Twinkle, EdU, and timeline of the experiment to determine the kinetics of mtDNA replication and proliferation of cumulus cell from meiosis arrest to meiosis resumption. Green, Twinkle; red, EdU; Right, quantification of EdU-positive and twinkle-positive cumulus cells from meiosis arrest to meiosis resumption, data from two biological replicate experiment. T-2^nd^, tiny secondary follicle; M-2^nd^, modest secondary follicle; P24h, PMSG 24h; P48h, PMSG 48h; H3h, HCG 3h. **(B)** Left, representative confocal image of the localization of FSHr, EdU, and timeline of the experiment to determine the kinetics of FSH capture capability and proliferation of cumulus cell from meiosis arrest to meiosis resumption. Green, FSHr; red, EdU; Right, quantification of EdU-positive and FSHr-positive cumulus cells from meiosis arrest to meiosis resumption, data from two biological replicate experiment. **(C)** Representative images of co-localization of FSHr and Twinkle. Hollow arrowheads indicate Twinkle, arrows indicate FSHr, solid arrowheads indicate co-localization of Twinkle and FSHr. Green, Twinkle; red, FSHr. **(D)** Representative images of Twinkle-positive cumulus cells stretched thick TZTs to connect with oocytes. arrowheads indicate thick TZTs. **(E)** Representative images of Twinkle-positive cumulus cells in re-COCs. **(F)** Representative images of FSHr-positive cumulus cells in re-COCs. **(G)** Model for germline-soma transporting mitochondria into its oocytes. FSHr positive cells capture FSH secreted by hypothalamus, not only improve their own FSHr expression level to occupy more FSH for meiosis arrest and oocyte growth, but promote mtDNA helicase-twinkle expression and TZTs assemble in cumulus cells to transport mitochondria into the oocytes. The mitochondrial transport is gradually terminated by LH level elevation. Green, mitochondria; purple, twinkle; magenta, FSH and FSHr; black, LH and LHr.

Our data showed that germline-soma provide mitochondria for mtDNA accumulation and mouse oocyte development via TZTs. Combined with previous report, which found that cytoplasmic transfer assisted mouse germ cells in growing into the largest cells prior to primordial follicle formation *(27)*, we speculated that mtDNA inheritance in mouse germline may share similar mechanism with other species, of which precursor germ cells receive organelles and cytoplasm from neighboring nurse cells to grow and therefore become oocytes *(28-32)*. Thus, the germline-soma-oocyte transport of mitochondria gives rise to 10^5^ copies of mtDNA in growing oocytes without replicating 9-14 times of a subpopulation of genomes in primordial germ cells during mouse oogenesis. We further propose that the germline-soma-oocyte transport of mitochondria underlies the constant transmission of mammalian mtDNA from one generation to the next. Our findings demonstrated the mitochondrial biogenesis with improved protection and higher efficiency than previous proposed in mammalian oogenesis.

## Acknowledgments

The authors thank M. Jiang for confocal technical support, Y Wang for TEM sample prepare, Y. Yang for polishing the manuscript and E. Tang for drawing of pattern graph.

## Author contributions

H.S., Y.Z. and X.D. supervised and designed the experiments. H.S. manipulated COCs, spatiotemporal staining and regenerated re-COCs. Zhao Y., Zhen Y. and S.S. tracked mitochondrial transfer in living COCs and re-COCs. Zhao Y. labeled mtDNA with EdU staining. Zhao Y., Zhen Y. and J.P. quantified mtDNA copies. Zhen Y., S.S. and J.P. performed immunofluorescence and confocal. S.S. observed mitochondrial morphology under transmission electron microscopy. Zhen Y. and J.P. analyzed data. H.S. organized the figures. Y.Z. and H.S. wrote the manuscript. Y.Z. and X.D. reviewed and edited the manuscript.

## Data and materials availability

All data are available in the manuscript or the supplementary materials.

## Supplementary Materials

Materials and Methods

Figs. S1 to S12

Tables S1

Captions for Tables S2 to S8

References (*33*–*37*)

