## Supplementary file for "Germline-soma Supply Mitochondria for mtDNA Inheritance in Mouse Oogenesis"

**This PDF file includes:**

Materials and Methods

Fig. S1 to S12

Table S1

Captions for Table S2 to S8

References *(33-37)*

**Other Supplementary Materials for this manuscript include the following:**

Table S2 to S8

**Methods and materials**

**Animals and reagents**

**Animal**

All mice experiments performed were under NIH Guide for the Care and Use of Laboratory Animals and approved by the Laboratory Animal Service, Fudan University (201802191S). Female BDF1 mice generated in this study were F1 hybrid of C57BL/6 and DBA mice and were hosted up to five per cage in an SPF-grade environment (20-24℃, 60-70% relative humidity, 12 h light/12 h dark rhythm). Water and food were available *ad libitum*.

### Chemical reagents and Antibodies

Chemical reagents and antibodies used in this study were summarized in Table S1.

### Light Microscopy

High resolution, bright field and fluorescence microscopy was performed with Nikon Eclipse Ti2 Confocal Microscope, NIS-Element AR (Nikon, Japan) or NikonC2 Si Confocal Microscope, NIS-Element AR (Nikon, Japan).

### Transmission electron microscopy (TEM)

For TEM, samples were prepared following standard protocols. Ovary and COCs were fixed with 2.5% glutaraldehyde and postfixed with 1% osmium /1.5% potassium ferrocyanide, dehydrated in graded alcohol series. Then COCs were permeated in increasing concentrations of Epon and ultimately embedded in 100% Epon for 48 h at 60°C. Consecutive ultrathin sections (80nm) were contrasted with lead citrate and imaged in a Zeiss EM 10 at an accelerating voltage of 80 kV. Ultrathin sections of 70 nm thickness were cut using a Leica ECT microtome (Leica, Germany), placed onto a 200-mesh formar-coated copper grid and post-stained using 4% uranyl acetate and

Reynold's lead. Then samples were contrasted with uranyl acetate and examined at an accelerating voltage of 120 kV with a JSM-4000 electron microscope (JEOL, Japan)

**Collection of MII oocytes and Cumulus-oocyte complexes (COCs)**

21-day BDF1 mice were superovulated by consecutive injections of PMSG (2.5 IU) and hCG (2.5 IU) at 48 h apart. 15 h after hCG injection, cumulus-oocyte complexes (COCs) were released from oviducts into G-gamete. Cumulus cells were denuded by 3 min at incubation with 0.1% hyaluronidase (Sage IVF) for collecting MII oocytes. For collecting Cumulus-oocyte complexes at germinal vesicle (GV-COCs), 21-day BDF1 mice were superovulated by injections of PMSG (2.5 IU) 20 h. Following euthanasia, ovaries were removed from mice and transferred into handling medium. COCs were punctuated from ovarian follicles using 29G 1/2 insulin syringe needles. For collecting COCs at meiosis I (MI-COCs), 21-day BDF1 mice were superovulated by consecutive injections of PMSG (2.5 IU) and hCG (2.5 IU) at 48 h apart. 3 h after hCG injection, MI-COCs were punctuated from ovarian follicles using 29G 1/2 insulin syringe needles.

**Contactless co-culture of GV oocytes with cumulus cells *in vitro***

Denuded oocytes were obtained by removing the cumulus cells of GV-COCs using a mouth-controlled micropipette. Contactless co-culture between the denuded oocytes and the cumulus cells was conducted for 24 h in a defined pre-IVM medium, which consisted of 80% TCM199+ 0.0022 g/ml sodium pyruvate + 0.05 IU/ml FSH + 0.8 μg/ml estrogen + 20% FBS +10 μM melatonin +10 μM cilostamide *(33)*. After 24 h, pre-IVM medium were replaced with IVM medium, which included 80% TCM199+ 0.0022 g/mL sodium pyruvate + 0.075 IU /ml FSH+ 0.5 IU/ mL HCG+ 0.8 μg/mL estrogen + 20% FBS + 0μM melatonin and cultured for an additional 24 h.

**Spatiotemporal staining of natural COCs with Nonyl Acridine Orange (NAO)**

Spatiotemporal-staining (SS) method was established for tracking mitochondrial transfer between cumulus cells and oocytes. Briefly, COCs with multilayer of cumulus cells were stained with 25 ng/ml Nonyl Acridine Orange (NAO) for 8~11 min according to the thickness of cumulus cells (Fig. S2A). Subsequently, COCs were thoroughly rinsed for 75 min (15 min, 30 min, 30 min, successively) in G-gamete medium and were deposited into confocal dish for live-cell screening. Those COCs, in where NAO exclusively labeled cumulus cells without disturbing oocytes, were selected for next step (Fig. S4A).

***In vitro* culture of NAO-labeled natural COCs**

The NAO-labeled GV-COCs were cultured for 1 h and 5 h in a defined pre-IVM medium, which consisted of 80% TCM199+ 0.0022 g/ml sodium pyruvate + 0.05 IU/ml FSH + 0.8 μg/ml estrogen + 20% FBS +10 μM melatonin +10 μM cilostamide. Some of NAO-labeled GV-COCs were treated by 10 μM CBX (carbenoxolone disodium, CBX) as gap junction block group and followed 5 h culture in the pre-IVM medium (CBX-COCs) *(34)*. The NAO-labeled MI-COCs were cultured for 5 h in the pre0IVM medium as meiosis resumption group (MI-COCs group).

***De novo* assembly of TZTs to reconstruct COCs *in vitro***

COCs obtained from 21-day mice superovulated by injections of 2.5 IU PMSG were incubated in G-gamete medium and drawn in and out of a mouth-controlled micropipette to generate denuded oocytes and small clumps of cumulus cells. The oocytes and disaggregated cumulus cells were together deposited into a modified reconstructed medium afterwards to form reconstructed COCs (re-COCs) (Fig. S5), which consisted of 80% TCM199+ 0.0022 g/ml sodium pyruvate + 0.05 IU /ml FSH+ 0.8 μg/ml estrogen + 20% FBS +10 μM melatonin +10 μM cilostamide + 200~400 μg/ml oligo hyaluronic acid (2000~8000 Da). Following incubation for 28~30 h at 37℃, 5% $CO_2$, the cumulus cells-oocyte aggregates, namely re-COCs were individually isolated for detecting TZTs or next experiments (Fig. S8A). For tracking mitochondrial transfer, small clumps of cumulus cells were stained in the medium supplemented with 25 ng/ml NAO for 20 min and rinsed three times in G-gamete (15 min, 30 min, 30 min, successively). Then the stained cumulus cells and unstained oocytes (GV and MI) were mixed to produce NAO-labeled re-COCs for next experiment (Fig. S8B).

**Tracking NAO-labeled mitochondrial transfer**

***NAO quantification in the oocytes from natural COCs and re-COCs***

After culture *in vitro,* NAO-labeled natural COCs and re-COCs were drawn in and out of a mouth-controlled micropipette to denude cumulus cells. Afterwards, the oocytes were allotted to microdroplets for fluorescent quantification. The Z-series acquisition model was applied to obtain multilayer images starting from the maximum diameter of the oocytes with step at 1 μm and count for 11 layers. Stacked images were uniformly post-processed using Extended Depth of Focus (EDF) algorithm. Background fluorescent noise were normalized using an Offset parameter at -25.0. Then the regions of interest (ROIs) were drawn manually using a Bezier curve to portray the profile of oocytes. NAO intensity relative to ROIs was selected for quantification using NIS-Element AR (Nikon, Japan). Subsequently, each quantified GV oocyte was transferred into signal PCR tube with micropipette for mtDNA copies quantification.

***MtDNA copies quantification in the GV oocytes***

Single oocyte was lysed using Qiagen REPLI-g Single Cell Kit (Qiagen, Germany) to obtain total mtDNA in 10 μl volume as described *(35)*. MtDNA copies quantification of single oocyte was performed by Roche LightCycler 480 II (Roche, Switzerland) to a target template spanning nt1323–nt1447 of the MT-16S rRNA gene as described *(36)*. The primer sequences for qPCR were F5’-CTAGAAACCCCGAAACCAAA-3’, R5’-CCAGCTATCACCAAGCTCGT-3’. The exact standard product concentration was measured with Epoch Microplate Spectrophotometer (Biotek, USA) and serial dilutions of DNA varied from $10^2$-$10^8$ copies/μl were prepared using DNase-free $H_2O$. These standard gradients were employed to generate standard curve with co-efficiency of reaction $R^2 > 0.99$. Second-round PCR was employed for absolute quantification of copies within each denuded oocyte. The reaction mixture and cycling conditions were consistent with the first round. Unimodal distribution in melt curve analysis confirmed that only a single product of the expected Tm value was generated. The DNA templates of standards and oocyte lysates were co-analyzed to absolute quantify mtDNA copies in LightCycler® 480 Software, Version 1.5 (Roche, Switzerland).

***Calculation of △NAO intensity***

NAO intensity was plotted as piecewise functions of diameter by spline interpolation of zeroth order, defined as $f_1(d)$ for control group (COC 1h and co-cultured), $f_2(d)$ for experimental group (COC 5h and reconstructed), where *d* is diameter and equidistance:

$$d = d_{min} + \frac{(k-1)(d_{max} - d_{min})}{n} \quad (k = 1,2,3,\ldots,\mathrm{n})$$

The difference of NAO intensity (△NAO intensity) between two groups (COC 5 h versus 1 h, re-COCs versus co-cultured) was calculated:

$$\triangle \mathrm{NAO\ intensity} = f_2(d) - f_1(d)$$

***Calculation of △mtDNA copies***

MtDNA copies of control group (COC 1h and co-cultured) were plotted as piecewise functions of diameter by spline interpolation of zeroth order, defined as $g_1(d)$; mtDNA copies of experimental group (COC 5h and reconstructed) were defined as $g_2(d)$, where $d$ is diameter and equidistance:

$$d = d_{min} + \frac{(k-1)(d_{max} - d_{min})}{n} \quad (k = 1,2,3,\ldots,\text{n})$$

The difference of mtDNA copies (△mtDNA copies) between two groups (COC 5 h versus 1 h, re-COCs versus co-cultured) was calculated:

$$\triangle\text{mtDNA copies} = g_2(d) - g_1(d)$$

***<u>Correlation between △NAO intensity and △mtDNA copies</u>***

Correlation between two trends above was analyzed by linear regression. All analysis was performed in *Python* (*Python* 3.7.3; https://www.python.org).

**Immunofluorescence for whole-COCs, single oocyte and cumulus cells**

COCs, single oocyte and cumulus cells were fixed with 2% paraformaldehyde (PFA) in PBS for 20 min at room temperature in a dish with glass bottom. After fixation and washed three times in PBS with 1% BSA, permeabilization were carried out in 0.5% Triton X-100, 0.05% Tween in PBS (PBST) for 3 h, and then blocked in permeabilization buffer with 5% donkey serum (Jackson Lab) for 3 h. Incubation with primary antibody was carried out overnight at 4 °C. COCs, single oocyte and cumulus cells were washed 3 x 10 min in PBST. Incubation with the secondary antibody was carried out at room temperature for 3 h. Previous washes were repeated, and counterstain was carried out with 1 mg/ml Hoechst33342 (Invitrogen). Confocal image was visualized under Nikon Eclipse Ti2 Confocal Microscope. Primary and secondary antibodies used in this experiment are summarized in Table S1.

**EdU incorporation**

COCs, oocytes and cumulus from 21-D-old female mice were collected in M199 medium supplemented with 10% FBS, and were pretreated for 3 h at 37°C in 7 μM aphidicolin, an inhibitor of nuclear DNA polymerase, to block nuclear DNA synthesis without affecting mtDNA replication, as previously described *(32, 37)*. The block then was followed by a 3 h incubation in 10 μM EdU and 7 μM aphidicolin to detect mtDNA replication in pre-IVM medium, supplemented with meiotic inhibitor (10μM cilostamide) at 37℃, in 5% $CO_2$. For detecting cumulus cells proliferation, COCs were directly incubated in 10 μM EdU for 2 h. EdU detection was carried out using the baseclick EdU HTS kit (Sigma, Table S1) according to the user manual. Incubation with the primary antibody and secondary antibody was carried out as described above. Images were visualized under a confocal microscope with a 60X oil-immersion objective and signals in the Z-series multi-channel images were captured using Nikon Eclipse Ti2 Confocal Microscope (Nikon, Japan).

**Confocal image quantification**

EdU-positive, twinkle-positive and FSHr-positive cumulus cells were conducted by eye counting the number of cells stained with red and green fluorescence.

**Statical analysis**

All statistical analysis was performed using Prism 8. Unpaired Student's t-test was used to test differences of diameter, fluorescence intensity and mtDNA copies between two groups. The correlation between two variables (diameter, ΔNAO intensity and ΔmtDNA copies) was performed by Pearson correlation. P-value less than 0.05 was considered statistically significant.

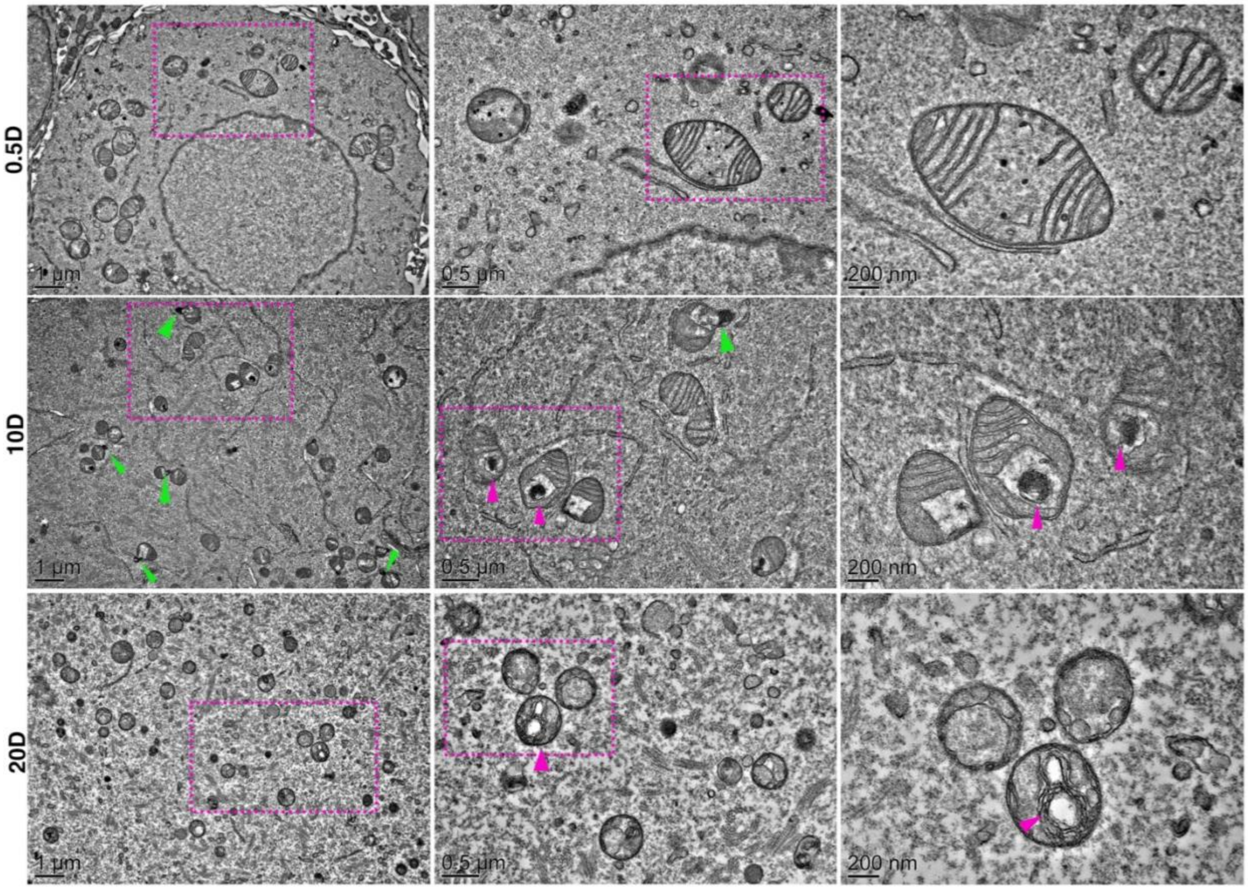


**Fig. S1. Elongated or dividing mitochondria were rare in the mouse oocytes at different developmental stages.**

There are round or oval mitochondria in mouse oocytes at 0.5day (upper), 10days (middle) and 20days (lower) of birth. Red arrowheads indicate mitophagy. For boxed areas, higher magnification images are shown. Green arrowheads indicate periphery fission of mitochondria.

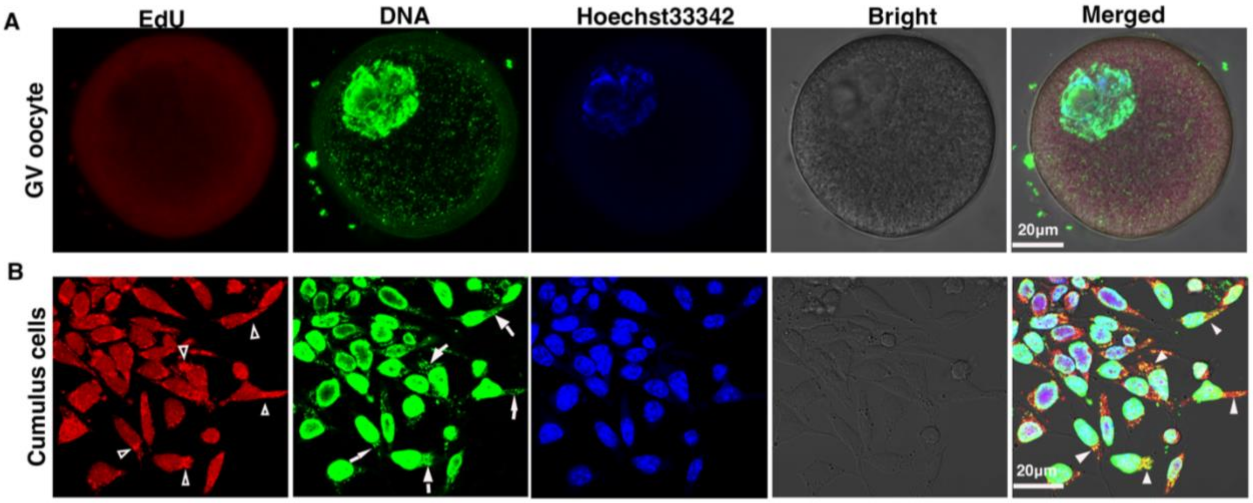


**Fig. S2. MtDNA replication was absent in mouse oocytes at germinal vesicle (GV).**

Representative images of GV oocyte (**A**) and cumulus cells (**B**) labeled with EdU (red) and anti-DNA antibody (green). Hollow arrowheads indicate EdU, arrows indicate mtDNA, solid arrowheads indicate co-location of EdU and mtDNA.

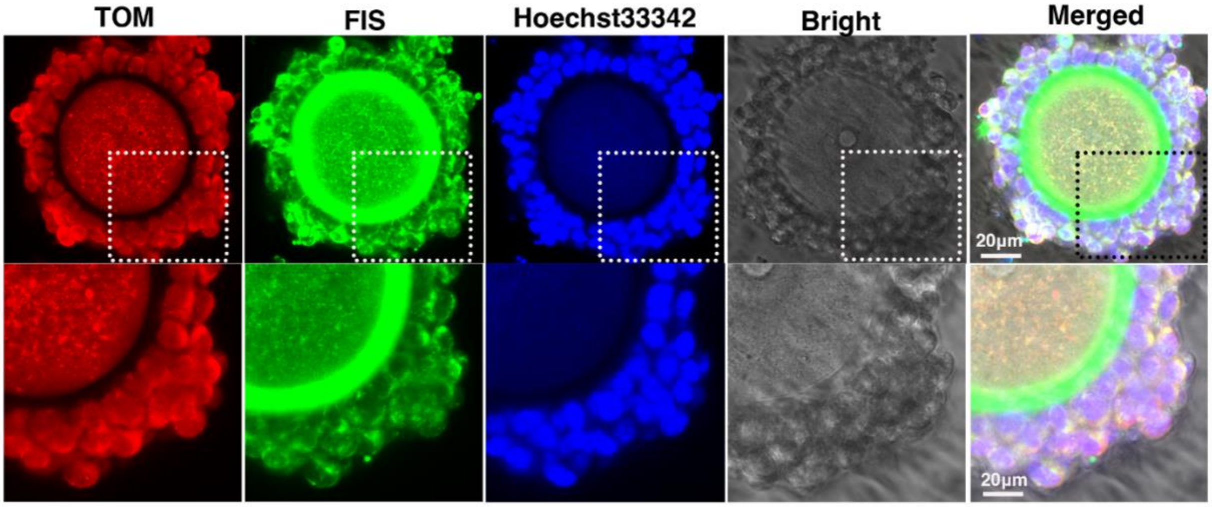


**Fig. S3. Peripheral division of mitochondria occurs in both cumulus cells and oocytes.** Periphery fission, which enables damaged material to be shed into smaller mitochondria destined for mitophagy, present in the cytoplasm of oocytes and cumulus cells.

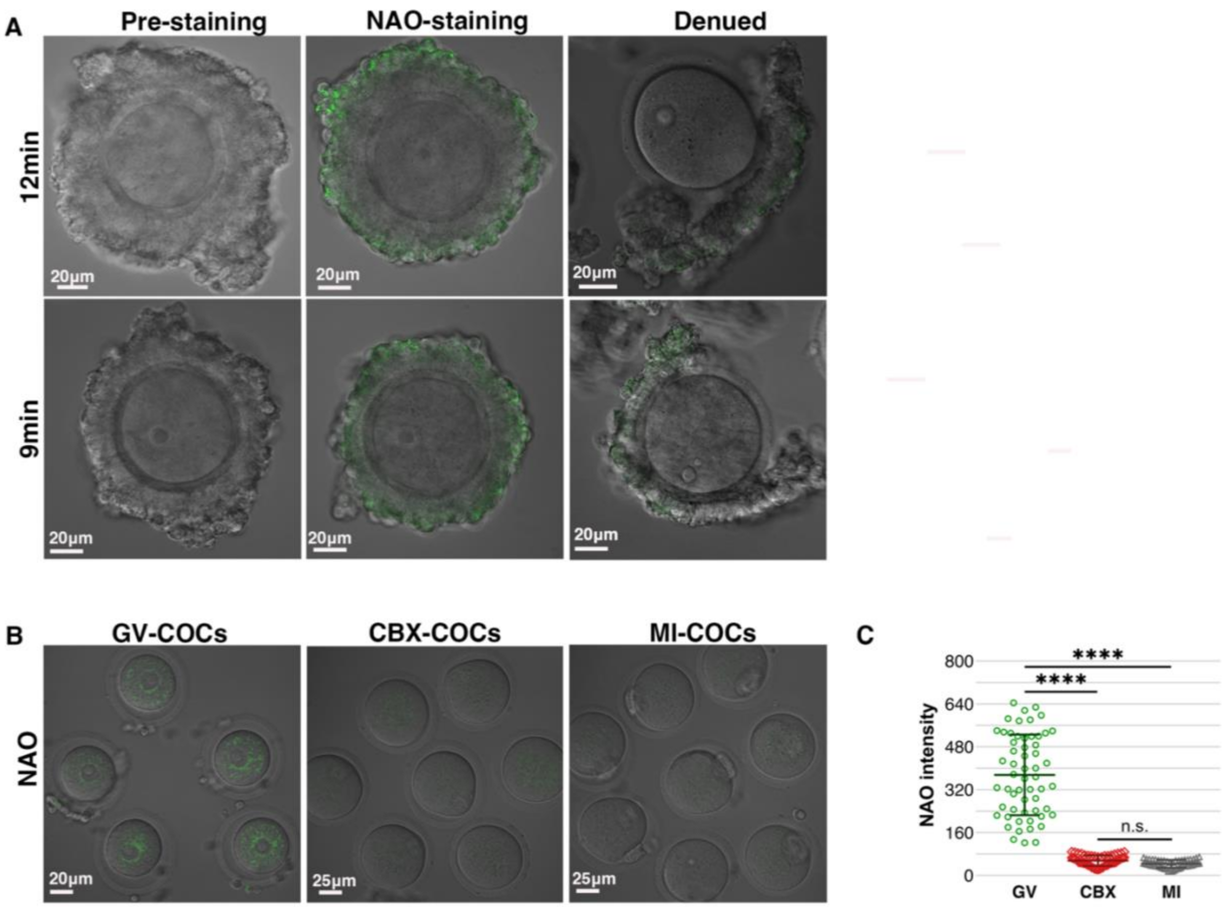


**Fig. S4. Tracking mitochondrial transfer between cumulus cells and oocytes by manipulating Spatiotemporal staining on mouse COCs with NAO.**

(**A**) GV-COCs with different thickness stained by NAO (left), by which as mitochondria probes only stain cumulus cells (middle) while the oocytes remained uncolored in COCs (right) by controlling staining time.

(**B**) NAO intensity in oocytes from three groups of COCs (GV-COCs, CBX-COCs, MI-COCs) after cultured 5 h *in vitro*.

(**C**) Quantification of NAO intensity in oocytes from the three groups COCs.

Background fluorescent noise in A, B and C was not normalized using an Offset parameter at -25.0. n.s., not significant, ****$P < 0.0001$.

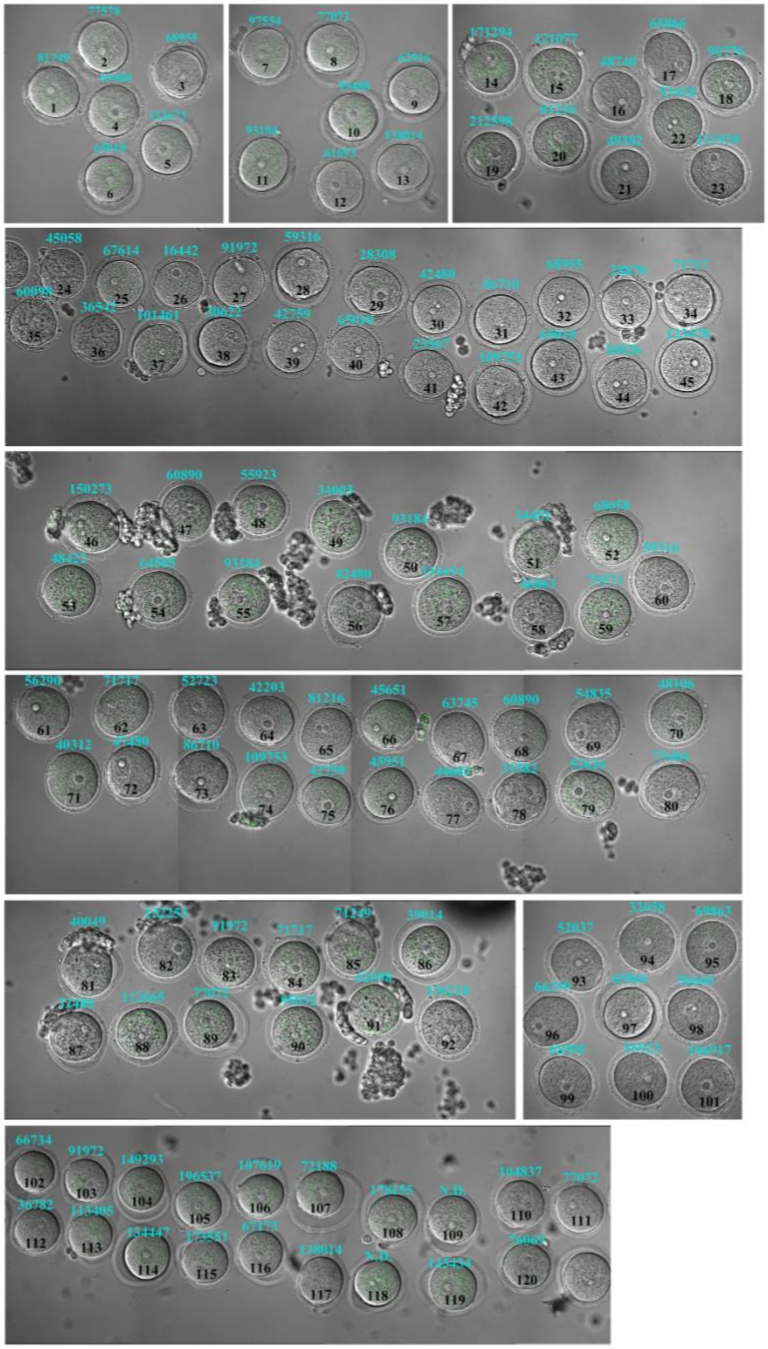

**Fig. S5. NAO intensity in mouse GV oocytes from COCs 1 h group (No.1 ~ 120).**

Corresponding mtDNA copies (cyan) and serial number (black) were consistently marked on oocytes. Background fluorescent noise was normalized using an Offset parameter at -25.0. N.D. represents not detected.

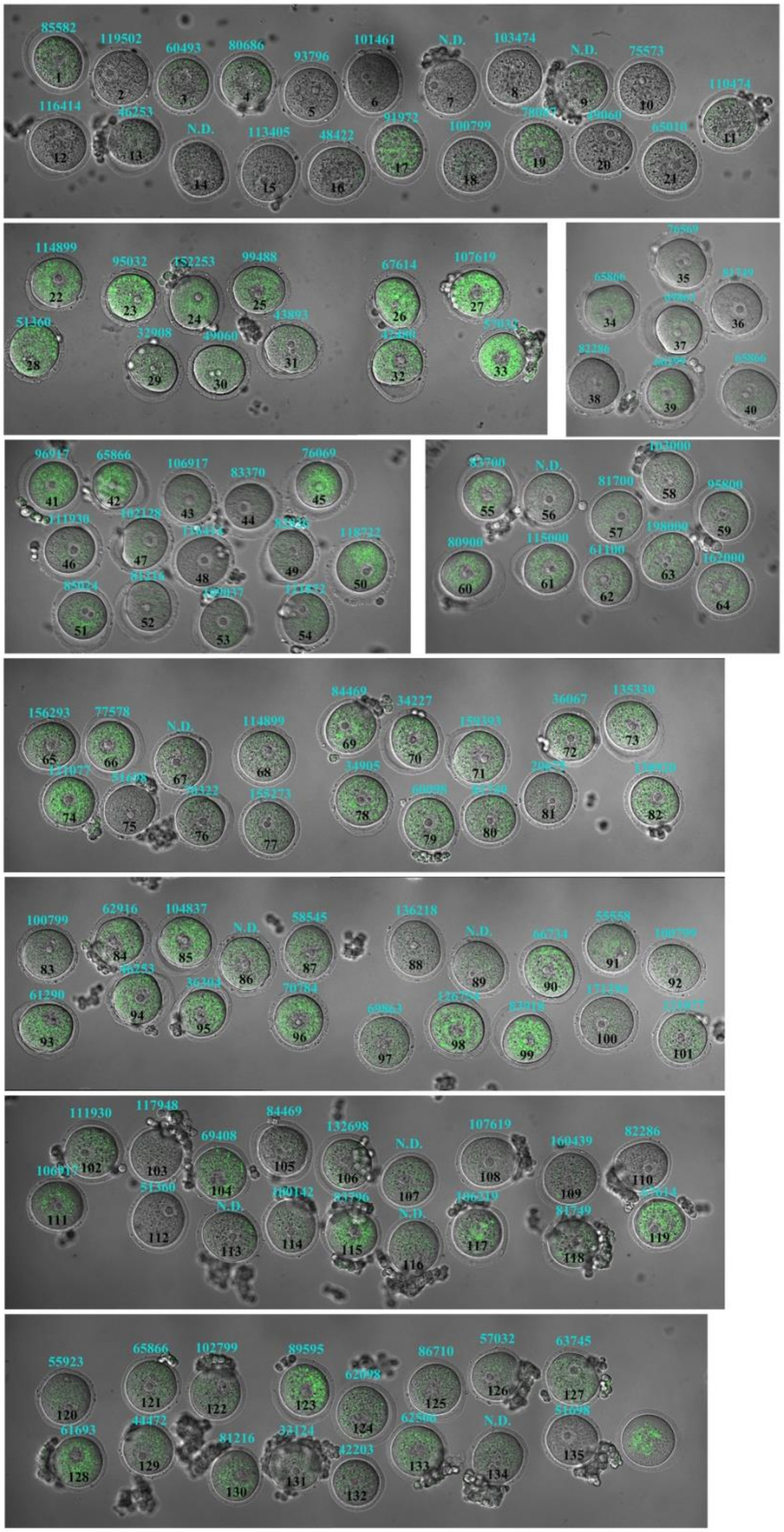
85582
119502
60493
80686
93796
101461
N.D.
103474
N.D.
75573
110474
116414
N.D.
113405
48422
91972
100799
65010
114899
95032
152253
99488
67614
107619
51360
32908
49060
43893
65866
82286
65866
96917
65866
106917
83370
76069
111930
102128
118722
83700
N.D.
81700
102000
95800
80900
115000
61100
198000
162000
156293
77578
N.D.
114899
84469
34227
36067
135330
34905
155273
100799
62916
104837
N.D.
58545
136218
N.D.
66734
55558
61290
36303
70784
111930
117948
69408
84469
132698
N.D.
107619
160439
82286
51360
N.D.
N.D.
81749
55923
65866
102799
89595
86710
57032
63745
62098
61693
44472
81216
62500
N.D.
51698
42203

**Fig. S6. NAO intensity in mouse GV oocytes from COCs 5 h group (No.1 ~ 135).**

Corresponding mtDNA copies (cyan) and serial number (black) were consistently marked on oocytes. Background fluorescent noise was normalized using an Offset parameter at -25.0. N.D. represents not detected.

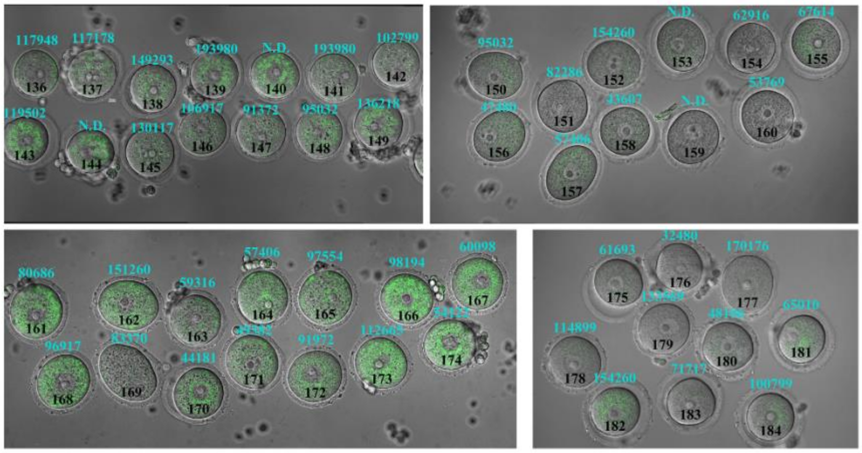


**Fig. S7. NAO intensity in mouse GV oocytes from COCs 5 h group (No.136 ~ 184).**

Corresponding mtDNA copies (cyan) and serial number (black) are consistently marked on oocytes. Background fluorescent noise was normalized using an Offset parameter at -25.0. N.D. represents not detected.

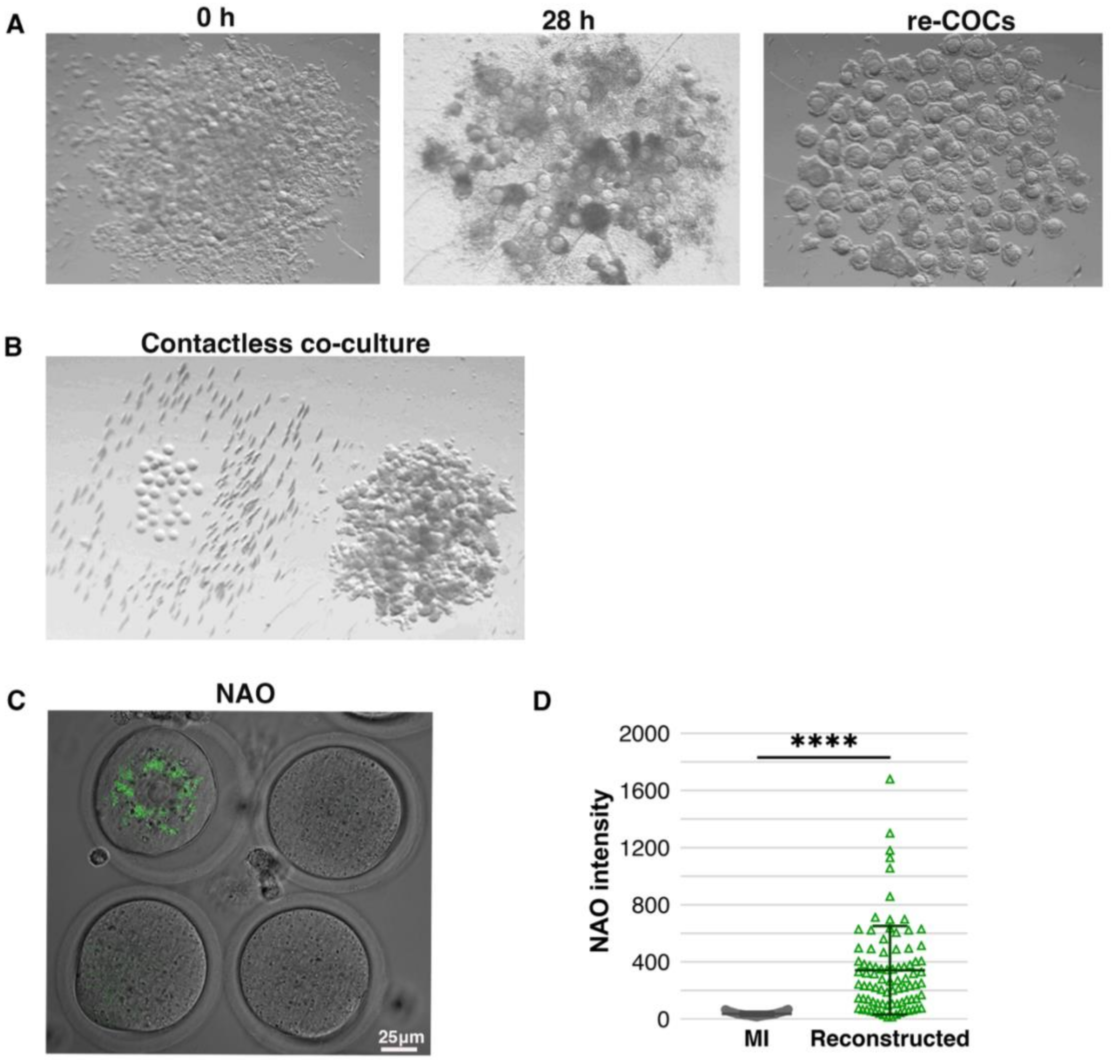


**Fig. S8. Assemble *de novo* TZTs for generating mouse re-COCs *in vitro*.**

(**A**) Representative images of re-COCs aggregation after mixing cumulus cells and GV and MI oocytes. Left, aggregating o h; middle, aggregating 28 h; right, re-COCs from middle picture.

(**B**) Representative images of co-culture of the GV oocytes during re-COCs generation.

(**C**) NAO intensity in oocytes cytoplasm of co-culture group and re-COCs group.

(**D**) Comparison of NAO intensity in GV and MI oocytes from re-GVCOCS and re-MICOCs. mouse oocytes of co-culture group and re-COCs group. Background fluorescent noise in C was normalized using an Offset parameter at -25.0. ****P < 0.0001.

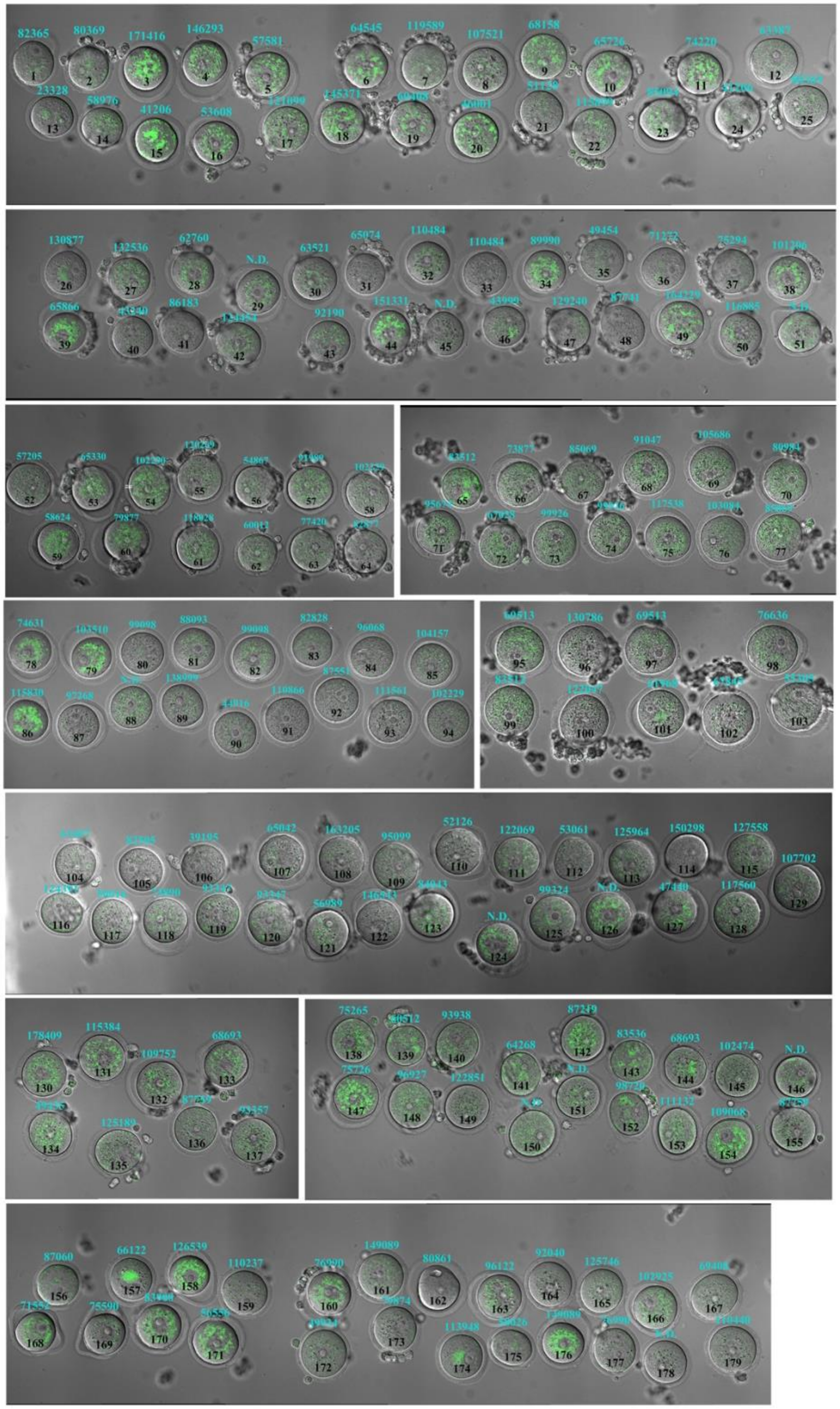

**Fig. S9. NAO intensity in mouse GV oocytes from re-COCs group (No.1 ~ 179).**

Corresponding mtDNA copies (cyan) and serial number (black) were consistently marked on oocytes. Background fluorescent noise was normalized using an Offset parameter at -25.0. N.D. represents not detected.

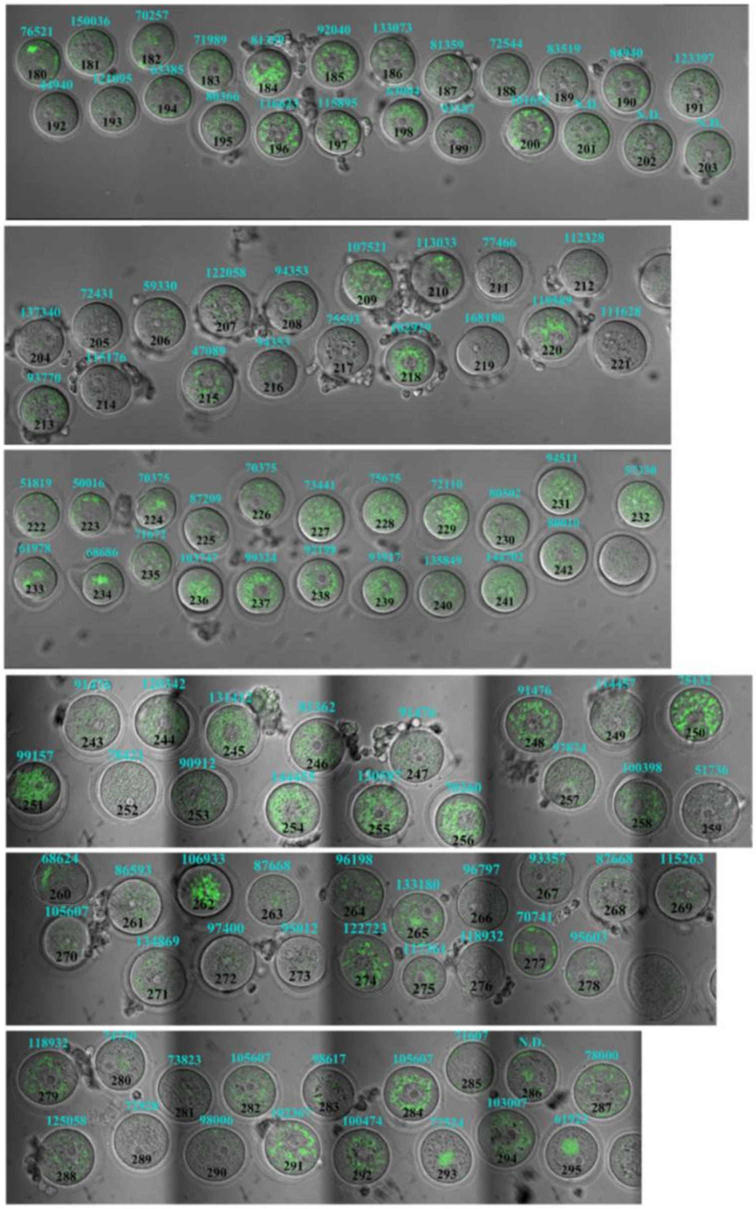

**Fig. S10. NAO intensity in mouse GV oocytes from re-COCs group (No.180 ~ 295).**

Corresponding mtDNA copies (cyan) and serial number (black) were consistently marked on oocytes. Background fluorescent noise was normalized using an Offset parameter at -25.0. N.D. represents not detected.

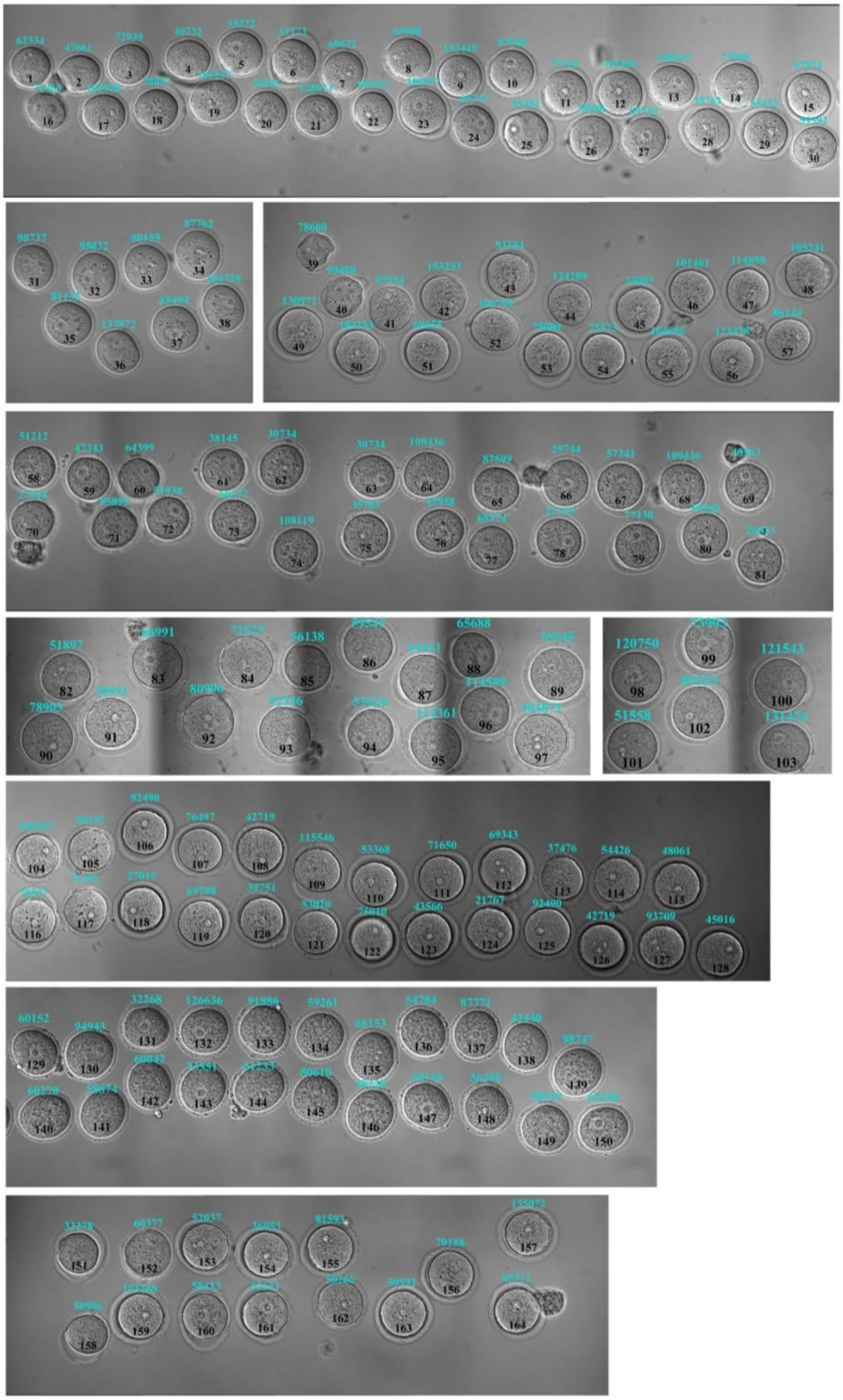

**Fig. S11. NAO intensity in mouse GV oocytes from co-cultured group (No.1 ~ 164).**

Corresponding mtDNA copies (cyan) and serial number (black) were consistently marked on oocytes. Background fluorescent noise was normalized using an Offset parameter at -25.0. N.D. represents not detected.

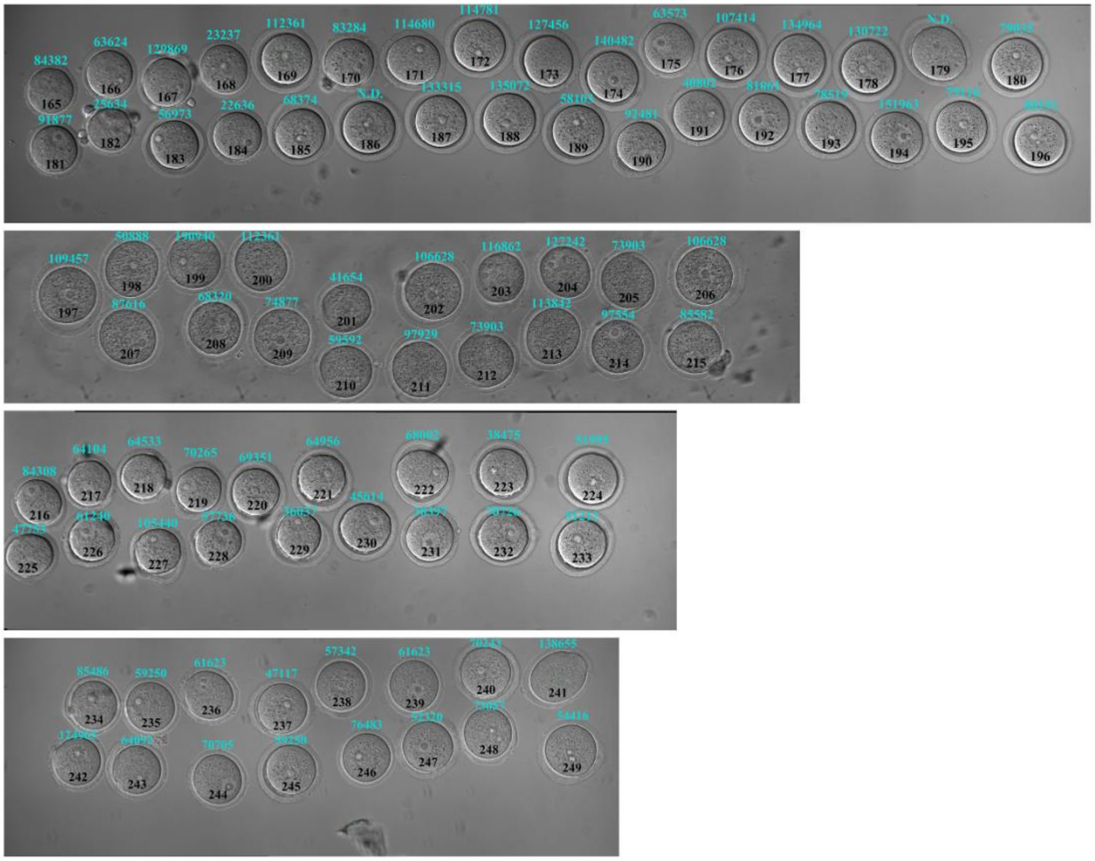

**Fig. S12. NAO intensity in mouse GV oocytes from co-cultured group (No.165 ~ 249).** Corresponding mtDNA copies (cyan) and serial number (black) were consistently marked on oocytes. Background fluorescent noise was normalized using an Offset parameter at -25.0. N.D. represents not detected.

**Table S1. Antibodies, chemicals and kits implemented in this study.**

| Reagent | | |
|---|---|---|
| **Antibodies** | **Vendor** | **Product number** |
| Rabbit Anti-TOM20 (1: 100) | Cell Signaling | 42406 |
| Mouse Anti-TOM20 (1: 100) | Sigma | WH0009804M1 |
| Rabbit Anti-MFF (1: 100) | Proteintech | 17090-1-AP |
| Rabbit Anti- FIS1 (1:100) | Abcam | ab229969 |
| Mouse Anti-Twinkle (1: 50) | Sigma | MABN814 |
| Rabbit Anti-FSHR (1:100) | Proteintech | 22665-1-AP |
| Mouse Anti-FSHR (1: 200) | Invitrogen | MA5-38525 |
| Alexa Fluor Plus 555 Phalloidin (1: 50) | Invitrogen | A30106 |
| Donkey Anti-Mouse IgG Alexa 488(1:500) | Jackson | 715-545-150 |
| Donkey Anti-Mouse IgG Alexa 594(1:500) | Jackson | 715-585-150 |
| Donkey Anti-Rabbit IgG Alexa 488(1:500) | Jackson | 711-545-152 |
| Donkey Anti-Rabbit IgG Alexa 594(1:500) | Jackson | 711-585-152 |
| **Culture medium and Chemical reagent** | | |
| TCM-199 | Sigma | M4530 |
| Pyruvic acid sodium | Sigma | P5280 |
| Estrogen | Sigma | E-2758 |
| FSH | Merck | N/A |
| Fetal Bovine Serum | Gibco | 12483020 |
| Triton™ X-100 | Sigma | X100 |
| Hyaluronic acid | Sigma | 40583 |
| PMSG | Ningbo Sansheng | 110254564 |
| hCG | Ningbo Sansheng | 110914564 |
| Nonyl Acridine Orange | Invitrogen | A1372 |
| Cilostamide | Sigma | C7971 |
| Carbenoxolone disodium | Sigma | C4790 |
| Phosphate Buffered Saline | Gibco | 10010-023 |
| Paraformaldehyde | Ted Pella Inc. | 18505 |
| Normal Donkey Serum | Jackson | 017-000-121 |
| G-gamete | Vitrolife AB | 10126 |
| G-IVF-plus | Vitrolife AB | 10136 |
| EdU Kit | Sigma | BCK-HTS594-2 |
| REPLI-g Single Cell Kit | Qiagen | 150345 |
| QuantiNova SYBR Green PCR Kit | Qiagen | 208056 |

**Table S2. Datasets for mtDNA copies of MII and oocytes *in vitro* maturation.**

**Table S3. Datasets for diameter measurement and NAO intensity quantification of oocytes from COCs 1 h and 5 h group.**

**Table S4. Datasets for diameter measurement and mtDNA copies quantification of oocytes from COCs 1 h and 5 h group.**

**Table S5. NAO intensity of oocytes from COCs at different stage.**

**Table S6. Datasets for diameter measurement and NAO intensity quantification of oocytes from co-cultured COCs group and re-COCs group.**

**Table S7. Datasets for diameter measurement and mtDNA copies quantification of oocytes from co-cultured COCs group and re-COCs group.**

**Table S8. NAO intensity of MI and GV oocytes after mixing cumulus cells with GV and MI oocytes 28 hours.**
